# 3D reconstruction of cell nuclei in a full *Drosophila* brain

**DOI:** 10.1101/2021.11.04.467197

**Authors:** Shang Mu, Szi-chieh Yu, Nicholas L. Turner, Claire E. McKellar, Sven Dorkenwald, Forrest Collman, Selden Koolman, Merlin Moore, Sarah Morejohn, Ben Silverman, Kyle Willie, Ryan Willie, Doug Bland, Austin Burke, Zoe Ashwood, Kyle Luther, Manuel Castro, Oluwaseun Ogedengbe, William Silversmith, Jingpeng Wu, Akhilesh Halageri, Thomas Macrina, Nico Kemnitz, Mala Murthy, H. Sebastian Seung

## Abstract

We reconstructed all cell nuclei in a 3D image of a *Drosophila* brain acquired by serial section electron microscopy (EM). The total number of nuclei is approximately 133,000, at least 87% of which belong to neurons. Neuronal nuclei vary from several hundred down to roughly 5 cubic micrometers. Glial nuclei can be even smaller. The optic lobes contain more than two times the number of cells than the central brain. Our nuclear reconstruction serves as a spatial map and index to the cells in a *Drosophila* brain.

## Introduction

The number of cells in a brain is a measure of its complexity, and has been used for comparing brains across species (Godfrey et al., 2021; Harrigan & Commons, 2015;Herculano-Houzel, 2011), developmental stages (Leuba & Kraftsik, 1994) and environmental conditions (Miller, 1995). Although *Drosophila melanogaster* is an important model organism, its brain cell number is still uncertain. Earlier accounts of the number of neurons in the adult fly brain (Alivisatos et al., 2012; Hsiao et al., 2016; Kaiser, 2015)(Chiang et al., 2011; Simpson,2009) were anecdotal. The first published estimates of *Drosophila* brain cell number appeared earlier this year. Godfrey et al. (2021) reported 88,300 cells. Raji & Potter (2021) reported 217,000 cells, of which 199,000 were neurons. These divergent estimates were produced by the same isotropic fractionator (IF) method (Herculano-Houzel & Lent, 2005). To verify the IF method, Godfrey et al. (2021) estimated 92,500 cells based on confocal imaging of an intact *Drosophila* brain.

We decided to count cells in a published 3D image of a *Drosophila* brain acquired by serial section electron microscopy (Zheng et al., 2018). As in the IF method, we assumed that nuclei are in one-to-one correspondence with cells. Nuclei in the 3D image were segmented via a computational pipeline consisting of three stages: a 2D convolutional network that classifies voxels as either nucleus or non-nucleus in all sections of the volume; a heuristic error detector and interpolator that examines continuity of the classification across adjacent sections and interpolates the classification at places of likely errors; and a final identification of connected components of nucleus voxels (Fig. 1A).

**Figure 1.**
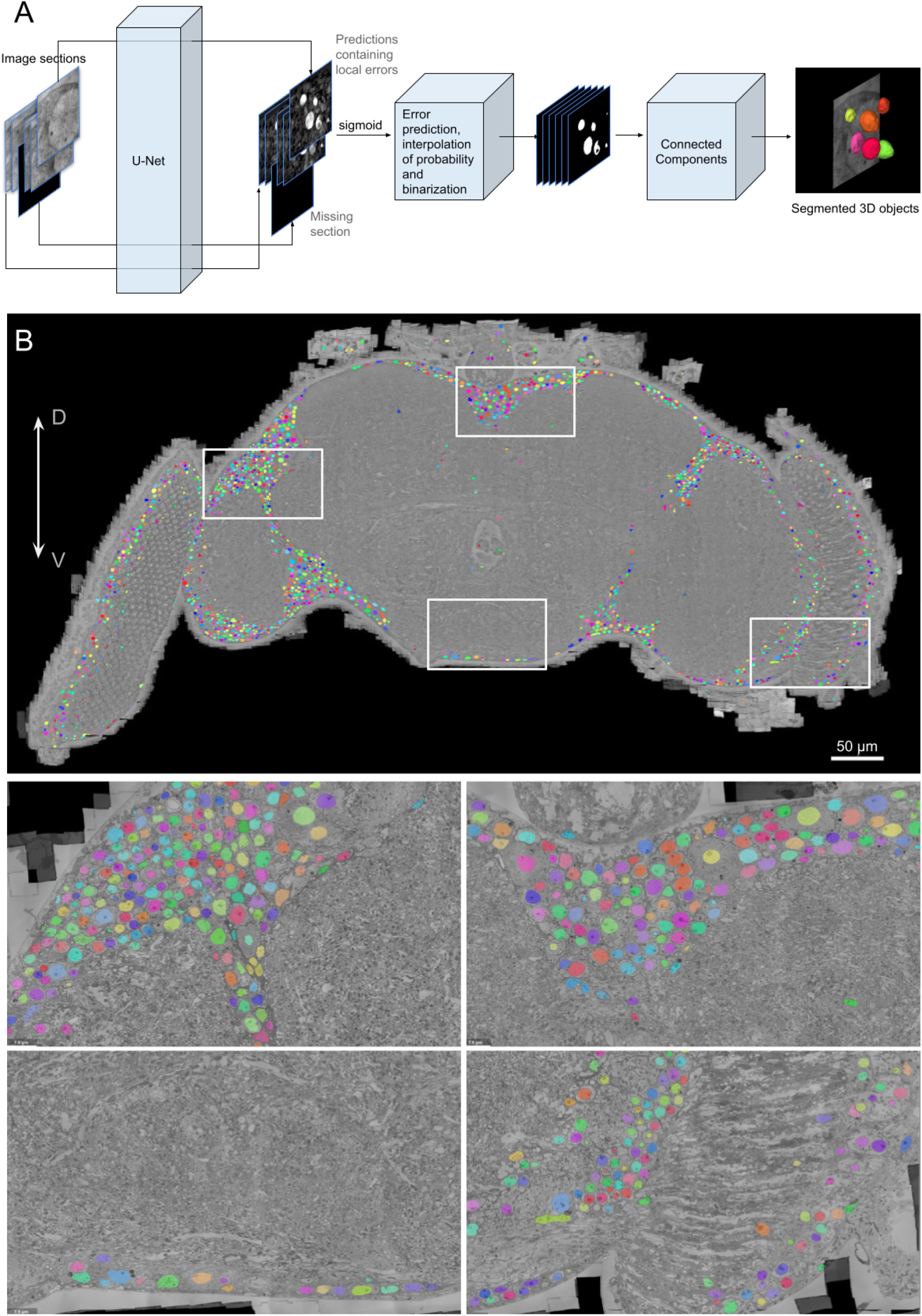
Segmentation of nuclei. **(A)** Computational pipeline translating raw input images into the final segmented 3D objects. A U-Net classifier sees image sections individually and for each pixel produces a prediction value, which after a sigmoid function can be seen as the pixel’s probability of belonging to a nucleus or not. A middle stage process detects likely errors in these probability values and replaces them with interpolated values from adjacent sections. A final stage process of connected components makes the final transformation from 0 and 1 voxel values into disjoint 3D objects. **(B)** Segmented nuclei (colored) from one (approximately transverse) section of the fly brain image volume. Bottom: zoom-in views of the boxed areas above. (D, dorsal; V, ventral.)

## Results

Segmented nuclei were located mostly at the borders of the central brain, optic lobes, and lamina (Fig. 1B). The interiors of these structures consisted of neuropil, defined as regions of entangled neurons and glia that are relatively free of cell bodies.

We examined thousands of image regions and found no false negatives. Out of a sample of 572 segmented objects (Methods), only a single false positive was large (12 μm^3^), and it occurred in a region outside the main brain tissue. All other false positives were smaller than 1 μm^3^.

Our count was potentially affected by other kinds of segmentation errors: merges of multiple nuclei into a single segment or splits of individual nuclei into multiple pieces. Such errors were primarily due to missing or misaligned sections in the underlying image volume. Some nuclei were outside the brain in partially or completely detached tissue nevertheless captured in the image volume. After controlling for all these factors (Methods), we estimate that there are 133,000 ± 3,000 nuclei in this fly brain. We further estimate that neurons make up at least 87% of this number (Methods), based on analysis of the above sample of 572 objects and their corresponding cell morphologies in FlyWire (Dorkenwald et al., 2020).

The largest nucleus in the brain has a size of 262 μm^3^. There are also nuclei as large as 450 μm^3^ in tissue surrounding the brain, for example in the antennal nerves. The smallest correctly segmented nuclei are about 3 μm^3^. Even smaller segmented objects tend to be false positives (not nuclei) and nucleus fragments mostly due to gaps and section misalignments in the image volume. In total, fewer than 100 nuclei in the brain are larger than 100 μm^3^ and 90% of all nuclei are smaller than 25 μm^3^ (Fig. 2A). Large nuclei are primarily concentrated around the frontal ventral face and the frontal midline of the central brain (Fig. 2B).

**Figure 2.**
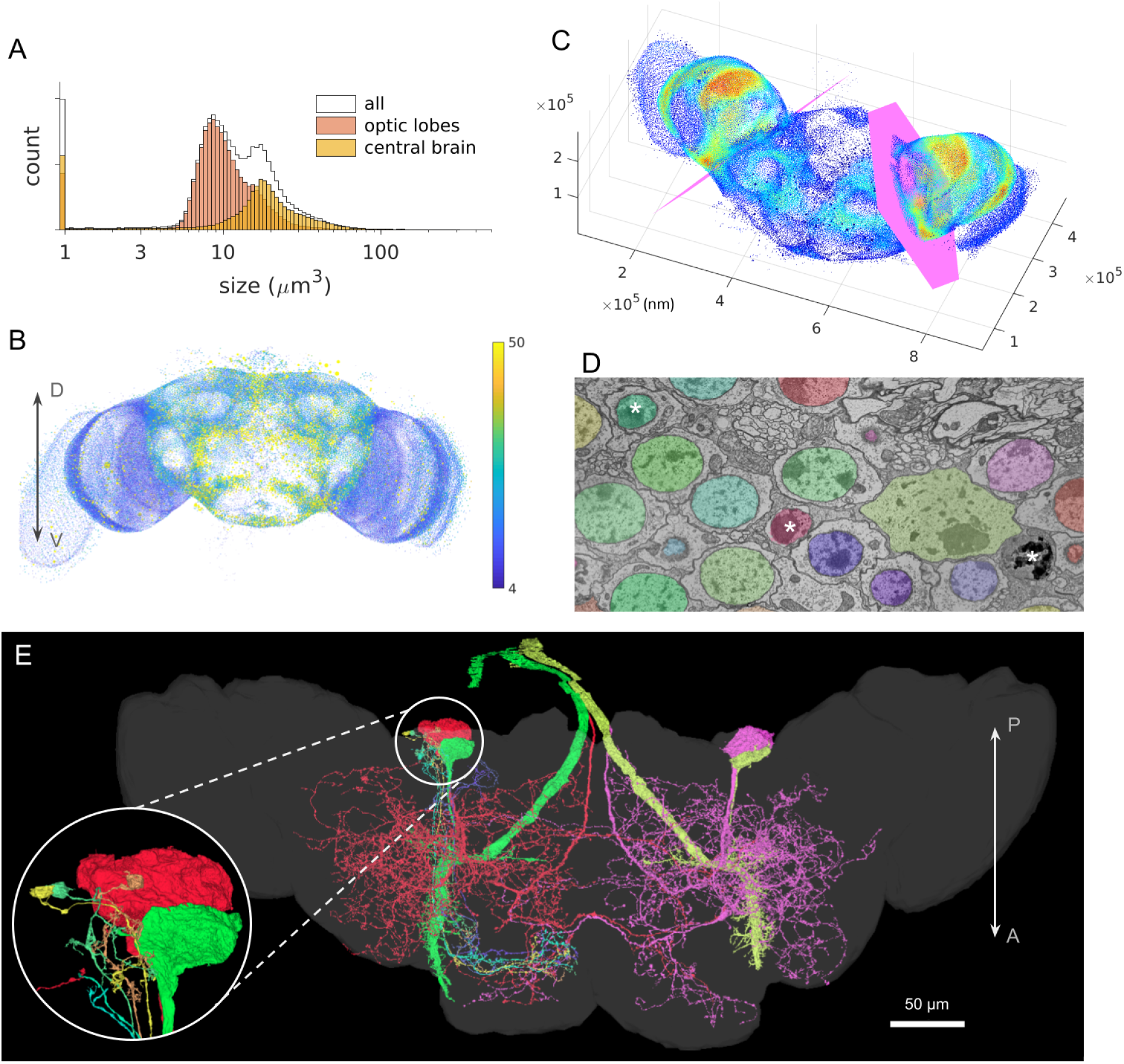
Statistics and quality evaluation. **(A)** Size distribution of all segmented objects. Segments in the smaller than 1 μm^3^ bin are false positives (not nuclei) and nucleus fragments mostly due to gaps in the image volume, commonly outside the main brain. **(B)** All segmented objects larger than 3 μm^3^ shown as a scatter plot of dots, size- and color-coded by volume (frontal view; dot sizes not to scale). **(C)** Scatter plot similar to B, of the same objects but color-coded by spatial density. Magenta planes denote boundaries used for the central brain versus optic lobe division in A. **(D)** Example region showing the segmentation on a single section. Note the three organelles marked by asterisks are visually similar but only two are true nuclei (colored). **(E)** A pair of neurons with the largest nuclei shown in red and magenta, the giant fiber cells in green and yellow, and a few small Kenyon cells in various other colors (dorsal view to show the relevant cell bodies, with the neuropil regions dimly shown for reference). Inset: tiny Kenyon cell somas contrasting the giant ones. (Axes: D, dorsal; V, ventral; A, anterior; P, posterior.)

As a final qualitative illustration and evaluation of the segmentation quality, we arbitrarily sampled a few Kenyon cells (known to be small neurons), again using FlyWire, and found their nuclei to be well segmented with volumes ranging from 16 to 32 μm^3^. The largest 2 neuronal nuclei (262 and 254 μm^3^ respectively) in the brain belonged to what appear to be the DNp32 descending neurons (Namiki et al., 2018), with their somas abutting the pair of giant fibre cell somas (at 208 and 183 μm^3^ in nucleus size; Fig. 2E). Most of the cells with the smallest nuclei around 3-5 μm^3^ were judged to be glial cells, but many other glial cells had regular-sized or huge nuclei.

We computed the spatial density of nuclei. The nuclei are most densely packed at the lateral and posterior edges of the optic lobes and along the anterior boundaries between the optic lobes and the central brain (Fig. 2C). This replicates what can be seen with nuclear staining in light microscopy (Fig. 7 in Robinow & White, 1991).

We assigned about 90,000 nuclei to the two optic lobes and 43,000 to the central brain, using two reference planes as approximate boundaries (Fig. 2C). The assignment is somewhat arbitrary, because the cell bodies at the borders between the optic lobes and central brain have ambiguous assignments (Fig. 1B). Shifting either reference plane by 10 μm can change the nucleus counts by as much as 4,000. Nuclei in the central brain are generally larger than those in the optic lobes (Fig. 2A,B).

## Discussion

Our estimate of *Drosophila* brain cell number is approximately equal (<3% difference) to the geometric mean of the two existing estimates based on the IF method (Godfrey et al., 2021; Raji & Potter, 2021). It is unclear why Raji & Potter (2021) report a number that is almost 2.5 times larger than that of Godfrey et al. (2021). We speculate that the large variation in nucleus volume (two orders of magnitude) could make it difficult to distinguish between intact nuclei and debris when counting with the IF method used by both studies.

Godfrey et al. (2021) reported an additional number based on confocal imaging of an intact brain, which was somewhat larger (<5%) than their IF number. Our number is still 44% larger than their confocal number. One uncertainty is that Godfrey et al. (2021) counted nuclei only in every third optical section, and interpolated for the other sections. Furthermore, it can be difficult to distinguish single nuclei versus overlapping close-by doublets in deep sections (Keating Godfrey, personal communication). Also, the amount of variation across individual flies is unknown, though perhaps it could be estimated from the variation of IF estimates.

Beyond the cell count, our nucleus reconstruction is a spatial map and index to the cells in a *Drosophila* brain, and can be used in a variety of ways. We have provided here a soma density map as one example. The nuclei are also being used by FlyWire, an online community for proofreading an automated reconstruction of the *Drosophila* connectome. Nucleus annotations can be used to detect neuron segmentation errors, as only single-nucleus segments have the potential to be correct. Soma location can be indicative of developmental origin or destination in neurogenesis (Ito et al., 2013; Yu et al., 2013). Relating nucleus and soma size may provide insight into cell function or metabolic needs (Willott et al., 1987; Yazdani et al., 2012). Somas of a specific cell or cell group can be used as anatomical landmarks (Wyman et al., 1984). The morphology of cell nuclei may be helpful for cell type classification (MICrONS Consortium et al., 2021).

We also expect that our methods and software will be useful for reconstruction of nuclei in the brains of other individuals and species, as more serial section electron microscopy datasets become available.

## Data availability

The nucleus segmentation can be viewed at the following URL: https://neuromancer-seung-import.appspot.com/?json_url= https://storage.googleapis.com/neuroglancer/drosophila_v0/nucleus/v5_z_intp_intp/seg/ng_state_ortho_slices.json

## Code availability

Code used to produce the nucleus segmentation and the data figures will be made available at https://github.com/seung-lab/fly-nuker

## Acknowledgements

We thank Zhihao Zheng for consulting on specific cells and for providing the original EM data, and Keating Godfrey for discussion on limitations in previous data.

## METHODS

### Segmentation of cell nuclei

We used an existing 3D image of an adult female *Drosophila* brain (Zheng et al., 2018), after the alignment of the serial section images was improved (Dorkenwald et al., 2020).

The segmentation algorithms consisted of three sequential processes: a U-Net classifier, a heuristic error detector and interpolator, and a final distributed connected-components grouping and stitching process.

We trained a standard convolutional 2D U-Net (with batch norms; (Ronneberger et al., 2015)) to classify individual image pixels as either nucleus or not nucleus. Human annotators manually annotated pixels in image sections, and about 600 512×512 pixel^2^ sized non-overlapping image sections at the 32×32 nm^2^ resolution from these annotations were fed to the U-Net to train the binary classifier. The human annotators had access to the adjacent image sections and the whole EM image volume while doing the annotation, and revisions were made whenever an error was discovered in these annotation labels. Additionally, heavy image augmentation was applied to the images during the training process, so the neural net would see different image statistics and object morphologies in its learning process. These augmentations included stretching and warping the images, altering the contrast and brightness, adding noise, as well as blanking out portions of the image with random values to simulate image defects often encountered in EM datasets. The trained U-Net was applied to the whole dataset as partially overlapping blocks using Chunkflow (Wu et al., 2021), generating a preliminary probability prediction of each voxel belonging to a nucleus or not.

Neither the EM volume nor the neural net is perfect. Because the trained neural net is a 2D and not 3D one, the prediction is independent across different and adjacent image sections, and image defects and prediction errors in any given section can therefore cause discontinuities in the nucleus prediction. Additionally, there are missing sections or swapped image section orders at certain locations in the dataset. Any such conditions could have caused a nucleus to become broken-up separate pieces in the final segmentation. We applied the top-hat transformations to the nucleus prediction values in the z direction (perpendicular to the sections) of the image volume to detect small-feature discontinuities, which is further cleaned up by binary morphological operations within-section to keep only those of significant sizes, and finally marked as likely errors. We refilled these locations, as well as known locations where the original EM image was blank (missing sections), with nucleus prediction values interpolated from adjacent sections. (The maximum allowed interpolated consecutive sections was 5, so a nucleus may remain broken in half if too many sections were missing.)

The previous interpolation step created the intermediary nucleus prediction map of the EM volume. Finally, a distributed connected components process (Turner et al., 2020) was applied to the prediction map to segment the nucleus-positive voxels into individual nuclei with independent numerical IDs. In this final step we preserved all connected components with a volume of at least 2000 voxels (0.08 μm^3^), resulting in a total of 143,140 segmented objects. Image section thickness was specified as 40nm, and a voxel size of 32×32×40 nm^3^ was assumed when converting voxel sizes into metric sizes.

### Validation of the nucleus segmentation

Human annotators were given 2020 8.1 μm × 8.1 μm 2D regions (systematically sampled by adding uniformly-distributed perturbation to evenly spaced grid points across the entire volume and only keeping those regions that are less than 90% blank), and tasked to find 1) if any of the nuclei in these regions were missed by the segmentation, and 2) if any of the segmented objects in these regions were not nuclei. 8 nuclei were found to have missed detection in these specific sections, while a total of 2587 segmented objects were present. However, all 8 cases had portions of their respective nuclei correctly detected on other image sections. Additionally, 6 false positive objects were encountered traversing these sections, all smaller than 1.5 μm^3^ in volume. Only 5 of these 14 error cases were from inside the brain as opposed to from partially detached tissue debris. An illustration of the nucleus detection quality can be seen in Fig. 2D, where true nuclei were segmented while mitochondria and nucleus-alike non-nucleus objects were not.

To assess the occurrence of non-nucleus false positives in terms of segmented 3D objects, we sorted all detected objects by size, and sampled every 250th for a total of 572 objects. As expected from the 2D nature of the artificial neural net and presence of section discontinuities in the image volume, except 1 fairly complete small nucleus outside the brain, all other 38 sampled objects smaller than 3 μm^3^ were found to be either false positives (15) or nucleus fragments, defined as pieces that are smaller than half of the imagined true size of the nucleus (23). For the objects larger than 3 μm^3^, 1 false positive was found in the image volume but was outside the brain. Together, these results show that our approach produces high quality nucleus segmentation where errors are few and small in size. Errors primarily occur at locations where image information is missing, or at severe image defects that occurred during stitching and alignment of image tiles into a 3D image stack.

To estimate the number of cells or nuclei in the brain, we needed to exclude those that came from partially or completely detached tissue that was clearly not part of the brain but were nevertheless captured in the image volume. We found 530 out of the 572 samples to be inside the main regions of the brain. 502 (87.8%± 1.4%, mean ± s.e.m of 4 equal-sized batches out of the 572 samples) were judged to be complete in-brain nuclei with minimal to no errors (482) or half-or-more portions of full in-brain nuclei (20) (Additionally, 438, or 87%, of these 502 confirmed in-brain nuclei were deemed unambiguously neurons, based on the morphology of their corresponding cell reconstructions in FlyWire). An additional 13 (2.3% ± 0.8%) were incorrectly merged double nuclei or triplets, encompassing 27 true nuclei in the brain. The true brain nuclei count from this sample of 572 objects were therefore 529. Extrapolation to the total number of 143,140 segmented objects (while accounting for double and triple nuclei) gave an initial estimate of 132,000 ± 3,000 nuclei in this fly brain.

We were aware of the presence of gross mergers due to image section misalignments in certain densely populated soma regions. Particularly, we additionally looked at all the top 200 objects in size (objects larger than 93 μm^3^). A total of 41 in-brain mergers in the top 200 were composed of roughly 500 nuclei. Adding these raises the total number of nuclei in this fly brain to 133,000 after rounding up. No false positives were encountered in the top 200.

